# *Toxoplasma gondii* disrupts vitamin B and K2 biosynthetic pathways in feline gut microbiota: microbial adaptation

**DOI:** 10.1101/2025.09.23.677982

**Authors:** Ji-Xin Zhao, Xuancheng Zhang, Shi-Chen Xie, Yi-Han Lv, Zhi Zheng, Ying-Qian Gao, Lu-Yao Tang, He Ma, Xiao-Xuan Zhang

## Abstract

Vitamins B and K_2_ (menaquinone) are critical for microbial and host processes, but their biosynthesis in feline gut microbiota and modulation by *Toxoplasma gondii* remain unclear. We assembled 1,553 high-quality feline intestinal bacterial genomes from metagenomic and public data to study vitamin B and K_2_ biosynthesis. We identified 86,829 genes and 159 enzymes, revealing regional differences in gene and species diversity for vitamin production. Among these, 782 genomes supported de novo synthesis of at least one vitamin. *T. gondii* infection altered synthesis pathways: early infection boosted pyridoxine and riboflavin synthesis in the small intestine, while menadione synthesis declined in both intestines. *Actinomycetota* and *Campylobacterota* were key to menaquinone biosynthesis, with *T. gondii* disrupting canonical pathways and activating alternatives. These findings shed light on the complex interplay between host, parasite, and microbiota in shaping vitamin biosynthesis, offering new perspectives for maintaining feline gut health during parasitic infection.

## 1. Introduction

The microbiome plays a pivotal role in maintaining gut and overall host health by mediating essential functions, including nutrient metabolism, immune system modulation, strengthening the intestinal barrier, and protecting against pathogens (Gensollen *et al*. 2016; Heiman *et al*. 2016; Hrncir 2022; Suchodolski *et al*. 2015; Suchodolski *et al*. 2012; Young *et al*. 2016). Among these functions, the synthesis of essential nutrients, such as vitamins, is crucial for supporting host health. While many vitamins are obtained from dietary sources, certain vitamins, including B_1_ (thiamine), B_2_ (riboflavin), B_3_ (niacin), B_5_ (pantothenic acid), B_6_ (pyridoxine), B_7_ (biotin), B_9_ (folic acid), and B_12_ (cobalamin), are also produced *de novo* by various bacterial species within the gut (LeBlanc *et al*. 2013a; Avior *et al*. 2015). Additionally, gut bacteria contribute to the synthesis of menaquinone, commonly known as vitamin K_2_, through the menaquinone biosynthetic pathway, which is mediated by enzymes such as MenA to MenG (Young et al. 2016). Microbial vitamin synthesis underscores the microbiome’s indispensable role in nutrient metabolism and its impact on host health.

Vitamins B and K are essential for processes such as energy production, cellular repair, amino acid synthesis, blood coagulation, bone health, cardiovascular function, and neurological function (Gallop *et al*. 1980; Strieker *et al*. 1996; Verbrugghe *et al*. 2013). Disruptions in these vitamin biosynthesis pathways, caused by factors such as dysbiosis, malabsorption due to enteritis (Edwards *et al*. 1987), inflammatory bowel disease, pancreatic insufficiency or hepatic lipidosis (Dutta *et al*. 1982; Bugrov *et al*. 2022; Center *et al*. 2000), dietary changes (Young et al. 2016; Bermingham *et al*. 2013a; Bermingham *et al*. 2013b), antibiotic treatments (Edwards et al. 1987), or parasitic infections (Duarte *et al*. 2016), can result in vitamin deficiencies. Such deficiencies impair vitamin-dependent processes, adversely affecting blood clotting (Strieker et al. 1996), bone health, immune regulation, and responses to infection (Scott *et al*. 2016).

Despite extensive studies on the role of the gut microbiome in nutrient synthesis in other species, such as humans and ruminants, the vitamin biosynthetic capabilities of the feline gut microbiome remain underexplored (Young et al. 2016). Existing studies have shown that parasitic infections, such as *T. gondii*, can significantly alter the composition and function of the intestinal microbiota (Meng *et al*. 2023). Given that the gut microbes plays a key role in synthesizing essential vitamins for the host, such as B vitamins and vitamin K₂ (Jiang *et al*. 2022; LeBlanc *et al*. 2013b), we speculate that *Toxoplasma* infection may disrupt these biosynthetic pathways, thereby impacting the host’s nutrition and health. *T. gondii* infections are common in felines, which serve as its definitive host (Elsheikha *et al*. 2021; Frenkel 1970; Hutchison 1970). During the sexual reproductive stage of *T. gondii* in the feline intestine, significant changes occur in the gut environment, affecting the microbiome’s composition, diversity, and functional capabilities, including its ability to synthesize vital nutrients (Bar *et al*. 2015; Bonnart *et al*. 2017). Understanding how *T. gondii* infections reshape the gut microbiome’s functional landscape can provide valuable insights into nutrient-host-microbe interactions, with implications for feline health and disease management.

While the effects of *T. gondii* on the host immunity and gut microbiota are documented (Hong *et al*. 2023; Lv *et al*. 2022), little is known about its influence on microbial vitamin biosynthesis in the feline intestine. *T. gondii* infections may alter the abundance and diversity of microbial species involved in vitamin B and menaquinone biosynthesis, disrupting key pathways (Benson *et al*. 2009). For instance, a decrease in *Bacteroides* populations may impair menaquinone synthesis, while shifts in *Lactobacillus* populations could impact vitamin B production. A deeper understanding of how *T. gondii* infection alters gut microbiota function, specifically vitamin biosynthesis pathways, could provide initial insights into its broader impact on feline health.

In this study, we collected a total of 7,324 feline gut bacterial genomes through metagenomic assembly and public data retrieval. Functional annotation of these genomes identified genes involved in the biosynthesis of vitamins B and menaquinone (vitamin K_2_). Using these reference datasets, we characterized regional heterogeneity in vitamin B and K_2_ biosynthesis within the cat gut microbiome. In addition, we explored how *T. gondii* infection reshaped the microbial pathways of vitamins B and menadione synthesis and investigated the menadione synthesis system in the cat gut microbiome through genomic analysis. The research provides new insights into host-parasite-microbiome interactions and their impact on nutrient synthesis, offering potential areas for future research in managing toxoplasmosis.

## 2. Methods

### 2.1 Experimental design and sample collection

Four litters (*n* = 18) of 4–5-month-old Chinese Lihua breed domestic cats (*Felis catus*) were obtained from local breeders. The cats were housed in a controlled environment and provided with commercial cat food, tailored to meet their daily energy requirements. Fresh water was made freely available to all cats. After one week of acclimatization, all cats underwent serological testing to ensure they were free from *Toxoplasma gondii* and other viral infections, including feline calicivirus, feline parvovirus, feline immunodeficiency virus, and feline leukemia virus, as outlined in previous studies (Acute *T. gondii* infection in cats induced tissue-specific transcriptional response dominated by immune signatures). Following the confirmation of their health status, the cats were randomly assigned into three experimental groups (*n*=6 per group): a control group (CON, prior to infection), asexual reproductive stage group (D3, third day of infection), and sexual reproductive stage group (D8, eighth day of infection, with oocyst excretion confirmed by microscopic examination of stool) (Frenkel *et al*. 1970; Shen *et al*. 2023).

*Toxoplasma gondii* PRU strain (Genotype II), commonly associated with human infections, was used for this study. The strain was maintained in laboratory Kunming mice. *Toxoplasma gondii* cysts were quantified using a light microscope and adjusted to 800 cysts/5 ml in PBS. Each cat in the experimental groups was orally infected with 800 brain cysts, while the control group received PBS orally. Prior to infection, as well as on the third and eighth days post-infection, all cats were euthanized via an overdose of isoflurane, administered by a qualified veterinarian with expertise in anesthetic procedures. Euthanasia was confirmed by complete unresponsiveness to stimuli and cessation of chest and heart movement. At least 2 g of small intestinal contents (comprising the duodenum, jejunum, and Ileum) and cecal contents were collected from each cat under sterile conditions and stored at -80°C for subsequent DNA extraction. Given that the microbial composition of the cecum closely resembles that of the colon in cats, the cecal contents were used as a representative sample of the large intestinal microbiota in this study (Hooda *et al*. 2012; Donaldson *et al*. 2016b).

All animal experimental procedures were approved by the Institutional Animal Care and Use Committee of Shanxi Agricultural University (Approval No. SXAU-EAW-2021XM121001). All animal experimental procedures were in compliance with the Biosafety Law of the People’s Republic of China.

### 2.2 DNA extraction and sequencing

Genomic DNA was extracted from small intestinal and cecal contents using the TIANGEN Magnetic Soil and Fecal DNA Kit, following the manufacturer’s instructions. DNA integrity was verified by 1% agarose gel electrophoresis, and genomic DNA concentration was quantified using the Qubit 3.0 Fluorometer (Invitrogen, USA) with the Qubit DNA Assay Kit. A total of 0.2 μg of DNA per sample was used as input for DNA library preparation. Sequencing libraries were constructed using the NEBNext® Ultra™ DNA Library Prep Kit (NEB, USA), in accordance with the manufacturer’s protocol. Index barcodes were incorporated into each sample for identification. Briefly, genomic DNA was fragmented to an insert size of 350 bp via sonication, and shotgun metagenomic sequencing was performed on the Illumina platform.

### 2.3 Public data collection

A total of 6,622 intestinal Metagenome-Assembled Genomes (MAGs) from domestic cats (*Felis catus*) were retrieved from the L’Entrepôt Pluridisciplinaire Recherche Data Gouv database. These genomes were derived from six public metagenomic projects (PRJNA758898, PRJEB9357, PRJEB4391, PRJNA944553, PRJNA908260, PRJNA923753), which collectively include 179 metagenomic samples from 102 cats. Additionally, we accessed a public project (PRJNA961122) hosted on NCBI, which includes 31 metagenomic samples from 23 cats.

### 2.4 Metagenomic assembly and binning

All bioinformatics tools and software used in this study were run with default parameters unless otherwise specified. A total of 67 metagenomic samples were included in the analysis. Illumina sequencing data were initially processed with FASTP (v0.23.0) (Chen *et al*. 2018) using the following parameters: “-q 20 -u 30 -n 5 -y -Y 30 -l 90 –trim_poly_g”. This step removed low-quality bases from the ends of reads and filtered out short or contaminated reads. To eliminate host contamination, Bowtie2 (v2.5.0) (Langdon 2015) was used to align the high-quality reads to the host genome (Genome GCF_018350175.1, NCBI), and host-derived reads were excluded. The remaining high-quality reads were used for subsequent analysis.

To generate comprehensive genomic data, each sample was assembled using MEGAHIT (v1.2.9) (Li *et al*. 2015) with the following optional parameters: “–k-list 21,41,61,81,101,121,141”. This resulted in 4,963,053 contigs, each with a minimum length of 200 bp and an average N50 length of 9,890.94 bp. Read mapping to these contigs was performed using BWA MEM (v0.7.17-r1188) (Li 2013), generating SAM files containing alignment information, which were then converted to BAM format with SAMtools (v.0.1.19-44428CD) (Li *et al*. 2009). Sequencing depths for the contigs were determined from the BAM files using the jgi_summarize_BAM_contig_depth script in the MetaBAT2 (v2.12.1) package (Kang *et al*. 2019).

Based on sequence features and sequencing depths, 3,610 contig bins (>2 kb) were generated using MetaBAT2 (v2.12.1) (Kang et al. 2019)with the following parameters: “-M 2000 -s 200000 –saveCls –unbinned”. These bins were further classified with SpecI (v.1.0) (Mende *et al*. 2013)and GTDB-tk (v1.1.1) (Chaumeil *et al*. 2019). Within each sample, bins with similar sequencing depth (±10%), G+C content (±0.02), and identical species assignments were merged. All bins were quality-assessed using CheckM2 (v1.0.1) (Chklovski *et al*. 2023), and those with completeness >50%, contamination <5%, and quality scores (calculated as: completeness – 5 × contamination) above 50 were retained. Redundancy was removed using dRep (v2.5.4) (Olm *et al*. 2017) with the options “cluster drep -pa 0.99 -nc 0.3 –SkipSecondary”, and bins with >99% ANI were grouped into one set. The highest-quality bin in each set, based on quality score, was retained as a Metagenome-Assembled Genome (MAG).

Open reading frames (ORFs) for these MAGs were predicted using Prodigal (v2.6.3) (Hyatt *et al*. 2010)with the option -p single. Clustering was then performed using the easy-cluster workflow in MMseqs2 (v7e284) (Steinegger *et al*. 2017) with the following parameters: --split-mode 2 --split-memory-limit 150G --cov-mode 2 -c 0.9 --min-seq-id 0.95 --cluster-mode 2 --cluster-reassign 1 --kmer-per-seq 200 --kmer-per-seq-scale 0.8. This process resulted in a non-redundant microbial gene catalog containing 1,575,958 genes.

### 2.5 Species-level clustering and phylogenetic analysis

To analyze species-level structure, MAGs with an ANI >99% were further clustered using dRep (v2.5.4) (Olm et al. 2017). MAGs with ANI >95% were considered to represent the same species. The MAGs with the highest quality scores were selected as representative species-level genome bins (SGBs), resulting in a total of 525 SGBs. Genome annotation was performed using Prokka (v1.14.5) (Seemann 2014), and phylogenetic trees were constructed at the amino acid level using PhyloPhlAn (v.1.0) (Segata *et al*. 2013). The resulting phylogenetic trees were annotated and visualized using iTOL.

### 2.6 Construction of a genome catalog of vitamin-related genes in the feline gut

Protein-coding genes from all MAGs were aligned against the Kyoto Encyclopedia of Genes and Genomes (KEGG) database(Kanehisa *et al*. 2000) using DIAMOND (v.0.9.22) (Buchfink *et al*. 2015)and BLASTP (Li *et al*. 2019)to obtain functional annotations for each gene. MAGs with the capacity for *de novo* biosynthesis of vitamins B and menaquinone were identified based on these annotations. An amino acid-level phylogenetic tree for these genomes was subsequently constructed using the method described earlier.

### 2.7 Mapping and quantification of microbial genomes in gut samples

High-quality sequencing reads (100 million reads) from 36 intestinal samples collected before and after *Toxoplasma gondii* infection were aligned to a microbial gene catalog using Bowtie2 (v2.5.0) (Langdon 2015)with maximal exact matching for optimal alignment. The resulting read counts were normalized to transcripts per million (TPM), and the relative abundances of the KEGG KOs were calculated based on gene abundance. Taxonomic profiles at the phylum and genus levels were determined by aggregating the abundances of genes assigned to each respective taxonomic category, with KO profiles derived in a similar manner. Functional roles associated with vitamin biosynthesis pathways were represented by KOs corresponding to specific functions, with the abundance of each functional role calculated by summing the relative abundances of the constituent KOs. To determine the pathway abundance for vitamins B and menaquinone, the abundances of all KOs involved in these biosynthetic pathways were aggregated.

### 2.8 Statistical analyses and visualization

Taxonomic and functional gene abundance data were used to calculate species richness and the Shannon diversity index. β-diversity was assessed via Principal Coordinate Analysis (PCoA) based on Bray-Curtis distance, and group differences were evaluated using permutational multivariate analysis of variance (PERMANOVA). The Wilcoxon rank-sum test was applied to assess the significance of differences in diversity indices, taxonomic units, and functional gene feature abundances across groups. Data normalization was performed using Min-Max scaling with the ‘caret’ package (v6.0.94). Rarefaction curves were generated using the ‘vegan’ package (v2.6-4), while a Sankey diagram was constructed using the ‘ggsankey’ package (v0.0.9). Network graphs were visualized using Gephi software (v0.10.1). All other visualizations were produced using the ‘ggplot2’ package (v4.2.3). All statistical analyses were conducted in R version 4.2.2.

## 3. Results

### 3.1 Genome assembly, gene catalog construction, and taxonomic profiling

This study represents a significant advancement in our understanding of the feline gut microbiome by generating a high-quality metagenomic assembly and microbial gene catalog. By integrating novel and publicly available datasets, we provide a comprehensive resource for exploring the taxonomy, diversity, and functional capacity of the feline gut microbial community.

To obtain a comprehensive feline gut metagenomic assembly, this study generated over 835.68 Gb of high-quality Illumina sequencing data from 67 feline gut and fecal metagenomic samples, including 36 samples from our study and 31 samples from public data, after quality control and host filtering. These metagenomic data were used for assembly and binning, and a total of 3610 raw bins were generated. Compatible bins with similar sequencing coverage (±10%) and G+C content (±2%) and the same taxonomic assignment were merged. A total of 702 MAGs were generated, with estimated completeness ≥70%, contamination <5%, and quality scores (defined as completeness-5×contamination) higher than 50. By combining 6,622 genomes from the L’entrepôt pluridisciplinaire Recherche Data Gouv database, a total of 7,324 feline intestinal genomes were collected in this study (Supplementary Table 1). After quality assessment (completeness ≥ 70%, contamination < 5%, and completeness - 5 × contamination ≥ 50) and clustering based on 99% average nucleotide identity (ANI), a total of 1,553 genomes met or exceeded the quality standards and were included in the subsequent analysis.

The MAGs were grouped into 525 species-level genome bins (SGBs) based on an average nucleotide identity (ANI) threshold of 95%, which is widely used as the species boundary for prokaryotic genomes (Figure 1A). The genome sizes ranged from 0.66 to 5.87 Mbp (average 2.31 Mbp), with GC content spanning 22.78% to 71.39% (average 44.59%). The number of ORFs per genome varied from 582 to 5,315 (average 2,176). The genomes had an average completeness of 94.94% and average contamination of 0.71%. Among the 525 genomes, 417 (79.43%) met high-quality standards, defined as having completeness ≥ 90% and contamination < 5% (Figure 1B-C). Taxonomic classification assigned all genomes to bacterial lineages, encompassing 16 phyla, 85 families, and 254 genera. The dominant phyla in the cat intestine included *Bacillota_A* (n=248, 47.24%), *Bacteroidota* (n=68, 12.95%), *Bacillota* (n=61, 11.62%), and *Actinomycetota* (n=61, 11.62%). At the genus level, the most abundant genera were *Collinsella* (n=16, 6.35%), Bacteroides (n=11, 2.10%), *Phocaeicola* (n=9, 1.71%), and *Blautia_A* (n=9, 1.71%).

**Figure 1.**
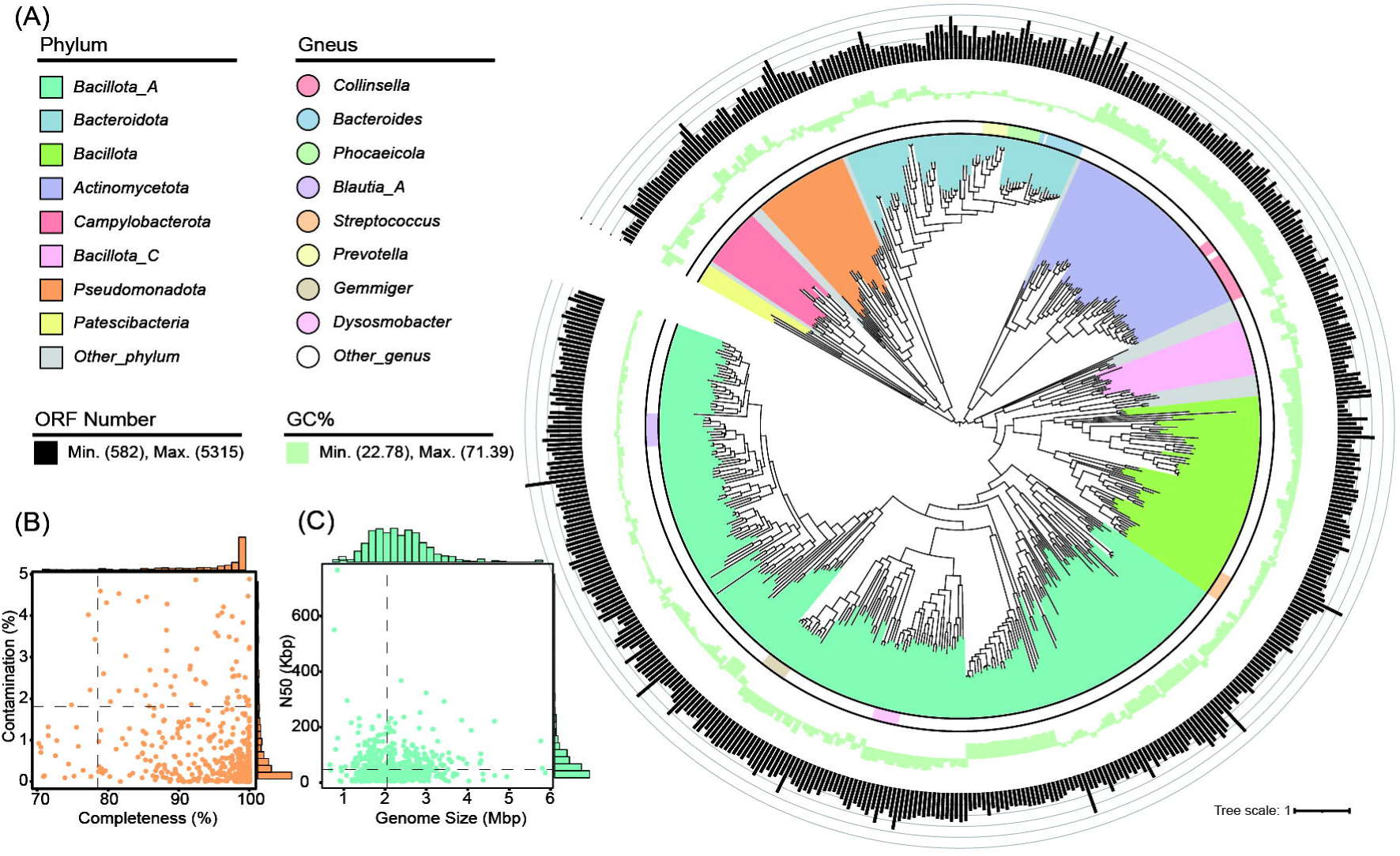
Overview of genomic and phylogenetic characteristics of 525 species-level genomes. **(A)** A phylogenetic tree displaying the relationships among 525 species-level genomes, with each clade color-coded according to its phylum-level classification. From inside to outside, the concentric rings represent: (i) genus-level classification, (ii) GC content of each genome (range: 22.78%–71.39%), and (iii) the number of open reading frames (ORFs) per genome (range: 582– 5315). **(B)** Scatter plot evaluating the completeness and contamination rates of the 525 genomes, showing high data quality across the dataset. **(C)** Relationship between genome size (in megabase pairs, Mbp) and N50 values (in kilobase pairs, Kbp), highlighting the assembly quality and size distribution of the genomes.

In addition, this study also constructed a non-redundant cat intestinal microbial gene catalog containing 1,575,958 genes with an average gene length of 886.38 bp. This genome and gene catalog are important tools for studying the classification and function of the cat gut microbiome.

### 3.2 Regional distribution of vitamin B and menaquinone biosynthetic genes

Analysis of the regional abundance, diversity, and functional pathways of vitamin B and menaquinone biosynthetic genes within the feline gut microbiome highlighted the interplay between microbial diversity, taxonomic composition, and metabolic specialization. To explore regional differences in vitamin biosynthesis gene abundance mediated by the feline gut microbiota, we compared the protein-coding genes of all MAGs with the KEGG database to obtain functional annotations for each gene. A total of 86,829 genes and 159 KEGG KOs were identified as involved in the synthesis of vitamins B (thiamine, riboflavin, niacin, pantothenic acid, pyridoxine, biotin, folic acid, and cobalamin) and menaquinone (Supplementary Table 2). As the sample size increased, the dilution curve gradually flattened, indicating that the KOs identified were sufficient to capture all genes associated with vitamin synthesis mediated by the cat intestinal microbiome (Figure 2A).

**Figure 2.**
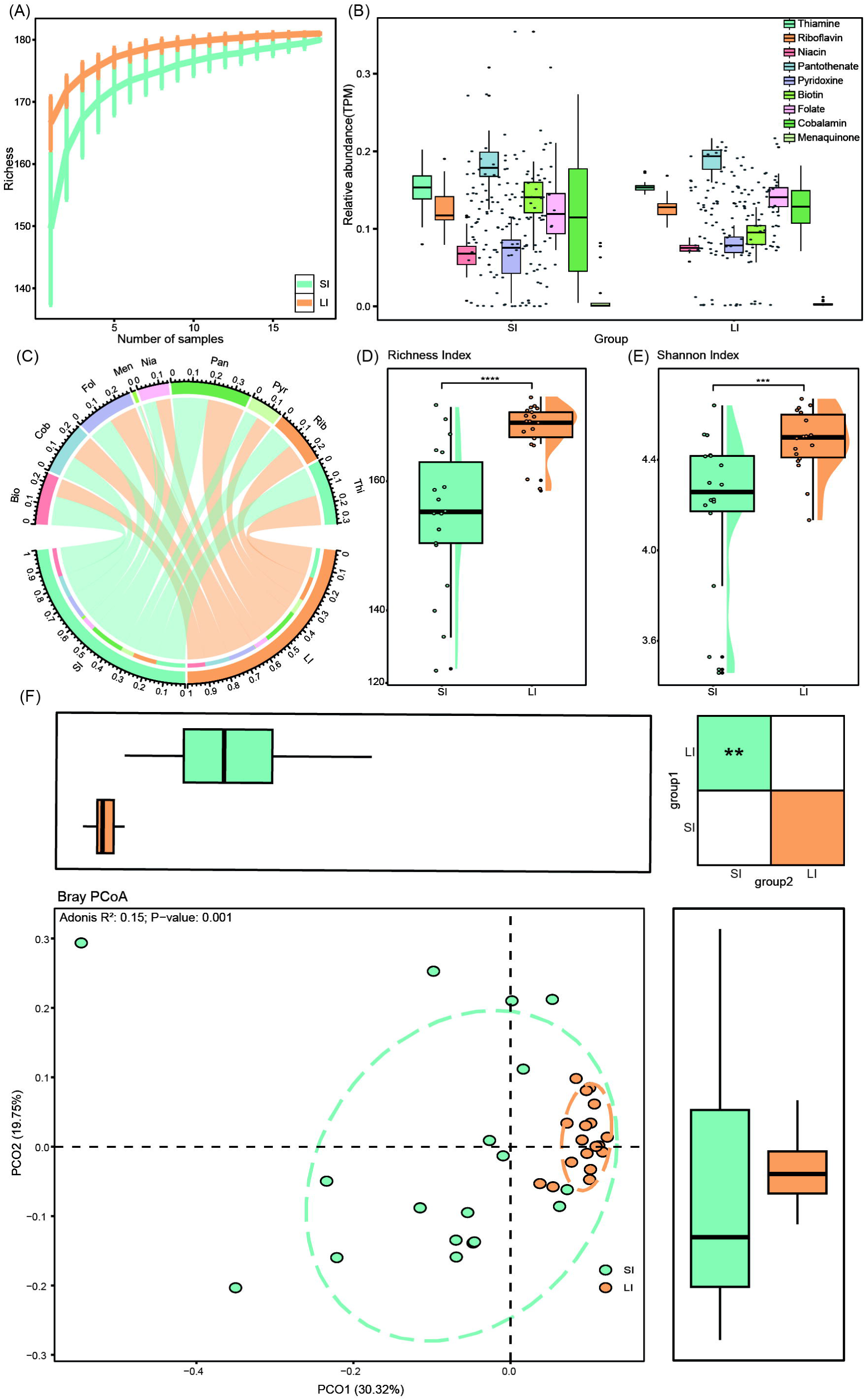
Microbial gene abundance, diversity, and community structure related to vitamin biosynthesis in the intestinal tract. **(A)** Rarefaction curve analysis illustrating the relationship between gene abundance and sample size, demonstrating the accumulation of gene diversity with increasing sample size. **(B)** Relative abundance of microbial genes associated with vitamin B and K_2_ biosynthesis in the large intestine (LI) and small intestine (SI). The horizontal line in each box represents the median, while the whiskers extend to the smallest and largest values within 1.5 times the interquartile range (IQR). **(C)** Chord plot showing the distribution of vitamin biosynthesis genes across different intestinal regions. Each region and vitamin is represented by a unique color. Vitamin B and K_2_ components are abbreviated as follows: Thiamine (Thi), Riboflavin (Rib), Niacin (Nia), Pantothenic acid (Pan), Pyridoxine (Pyr), Biotin (Bio), Folic acid (Fol), Cobalamin (Cob), and Menaquinone (Men). **(D-E)** Rain cloud plots, combining dot plots, box plots, and density plots, present the abundance and Shannon index of microbial genes involved in vitamin B and K_2_ biosynthesis. Dot plots show individual sample points, box plots indicate medians and quartiles, and density plots depict data distributions. Statistical significance was assessed using the Wilcoxon rank-sum test: \**p* < 0.05, \*\**p* < 0.01, \*\*\**p* < 0.001. **(F)** Bray-Curtis principal coordinate analysis (PCoA) illustrating microbial community structure across different intestinal regions. Ellipses indicate 95% confidence intervals. Boxplots above and to the right show sample scores along PCoA1 and PCoA2 axes, with medians and quartiles represented. Whiskers extend to the most extreme values within 1.5 × IQR. The heatmap in the top-right corner highlights significant differences between groups along PCoA1 and PCoA2 axes. Statistical significance was determined using the Wilcoxon rank-sum test (\**p* < 0.05, \*\**p* < 0.01, \*\*\**p* < 0.001).

Analysis of the relative abundance of genes involved in the synthesis of these nine vitamins in both the large and small intestines revealed that genes involved in thiamine synthesis were the most abundant, followed by those for riboflavin, folate, and cobalamin. In contrast, genes involved in menaquinone synthesis were the least abundant (Figure 2B-C).

To further investigate the microbial synthesis of vitamins in the feline intestine, we characterized the pathways involved in the biosynthesis of vitamins B and menaquinone across different intestinal segments. For thiamine, the biosynthetic pathway consists of two branches, with the thI branch being predominant in the cat’s intestinal microbiome. The biosynthesis of cobalamin involves both aerobic and anaerobic pathways, while the pathways for niacin, riboflavin, folic acid, and menadione are single-step. Interestingly, key genes in the folic acid biosynthesis pathway were found to be present at extremely low abundance in the feline intestine. Menadione synthesis occurs indirectly in the cat’s intestinal microbiome (Supplementary Figs. 1-3), suggesting that cats may rely on an external source of vitamin K_2_. Further analysis of gene diversity in the large and small intestines showed that the large intestine exhibited significantly higher diversity and richness of vitamin biosynthesis genes compared to the small intestine (*p* < 0.05) (Figure 2D-E).

PCoA of vitamin synthesis gene composition revealed distinct gene composition structures associated with vitamin synthesis in different intestinal regions, and PERMANOVA confirmed significant differences in these gene structures between the large and small intestines (R² = 0.14, *p* < 0.001). These findings indicate clear regional differences in vitamin synthesis within the feline intestine (Figure 2F).

### 3.3 Distribution and regional differences of vitamin biosynthetic genes

We investigated the microbial species distribution, diversity, and regional differences in vitamin biosynthetic genes within the feline gut microbiome, focusing on the role of specific taxa in synthesizing essential vitamins B and menaquinone across different intestinal regions. Rarefaction curve analysis showed that as the sample size increased, the cumulative sequencing data reached saturation, indicating that the sequencing effort adequately covered the microbial genome (Figure 3A). We assessed differences in species composition associated with vitamin synthesis by analyzing within-group and between-group relationships using alpha and beta diversity indices. Alpha diversity was calculated for each sample using the Shannon diversity and Richness indices. Comparative analysis was performed using Fisher’s method with p-values derived from the Wilcoxon rank-sum test. The results revealed that species diversity and richness in the large intestine were significantly higher than in the small intestine (*p* < 0.05) (Figure 3B-C). PCoA, based on Bray-Curtis distances, was used to evaluate species composition differences between the large and small intestines. Visualization of the first two principal axes explained 25.55% of the total variation, and significant differences in species composition were confirmed by PERMANOVA (R² = 0.1368, *p*= 0.001) (Figure 3D).

**Figure 3.**
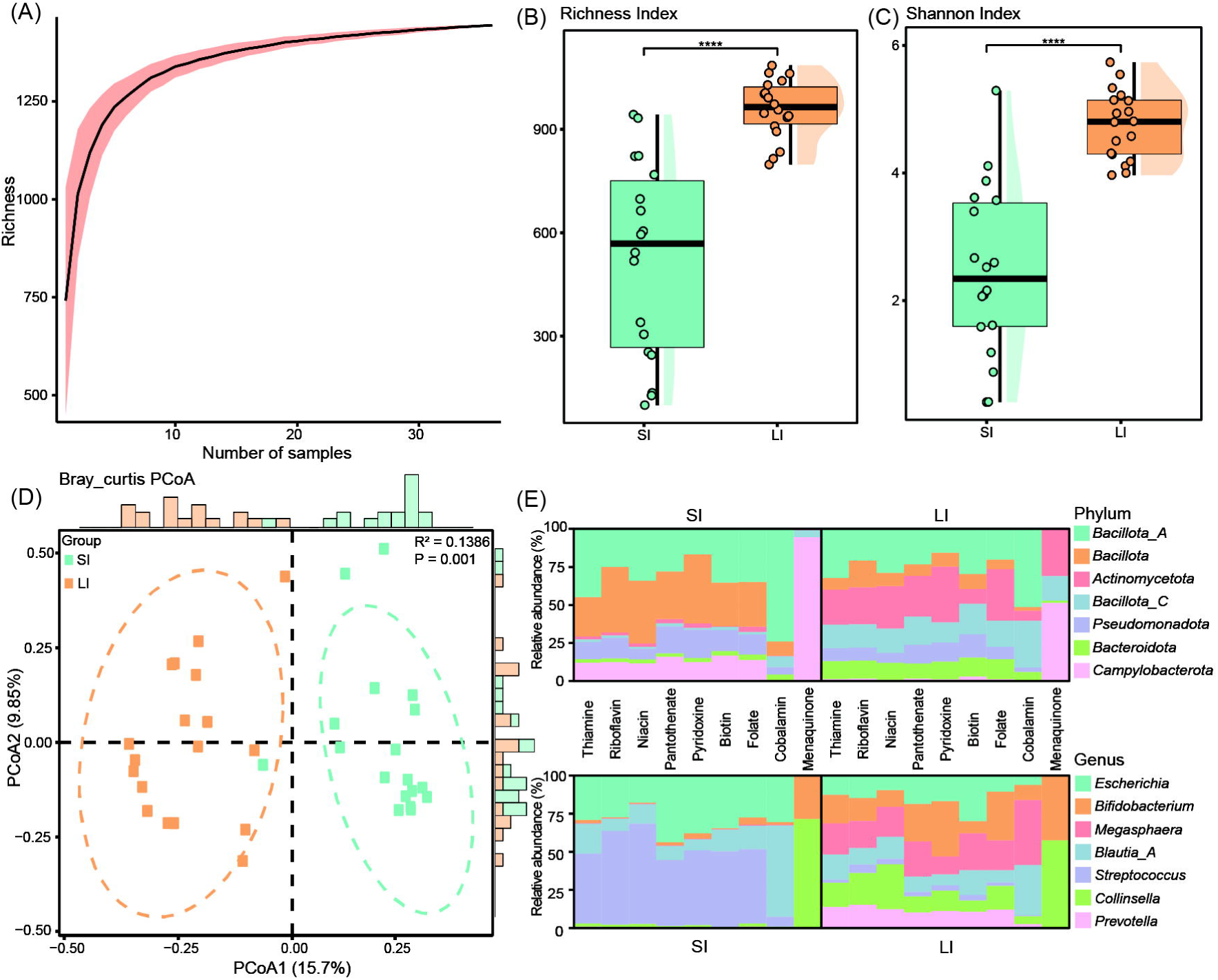
Microbial diversity and functional contributions to vitamin B and k_2_ biosynthesis. **(A)** Rarefaction curve illustrating the relationship between the accumulation of genome richness and increasing sample size. **(B-C)** Rain cloud plots integrating dot plots, boxplots, and distribution plots. The distribution plots depict the probability density of the data, the dot plots represent the distribution of individual sample points, and the boxplots summarize microbial genome richness and Shannon index values involved in vitamin B and K_2_ biosynthesis. Statistical significance was assessed using the Wilcoxon rank-sum test: \**p* < 0.05, \*\**p* < 0.01, \*\*\**p* < 0.001. **(D)** Scatter plot representing beta diversity and compositional changes of microbial genomes associated with vitamin B and K_2_ biosynthesis. Samples are plotted using the first and second principal coordinates (PCoA1 and PCoA2), with percentages of explained variance indicated. Ellipsoids denote 95% confidence intervals for each group. Marginal bar graphs above and to the right depict sample density distributions along PCoA1 and PCoA2. **(E)** Bar graphs showing the taxonomic distribution of gene sets related to vitamin B and K_2_ biosynthesis, categorized at the phylum and genus levels (top seven in abundance).

To identify the microorganisms involved in vitamin synthesis, we categorized the genes associated with microbial vitamin biosynthesis. At the phylum level, *Bacillota_A*, *Bacillota*, and *Actinomycetota* were the most abundant bacterial phyla associated with vitamin biosynthesis. Interestingly, *Bacillota* and *Bacillota_A* are primarily involved in synthesizing B vitamins, including thiamine, riboflavin, niacin, pantothenic acid, pyridoxine, biotin, folic acid, and cobalamin. The synthesis of menadione in both the large and small intestines was primarily carried out by *Campylobacterota* (Figure 3E and Supplementary Table 3). At the genus level, genera such as *Escherichia*, *Bifidobacterium*, *Megasphaera*, *Blautia_A*, *Streptococcus*, *Collinsella*, and *Prevotella* exhibited the highest abundance of genes involved in B and menaquinone biosynthesis (Figure 3E). While the relative abundance of *Escherichia* did not show significant differences between the large and small intestines, other genera exhibited significant changes in their relative abundance between the regions. The abundance of *Bifidobacterium*, *Megasphaera*, *Blautia_A*, *Collinsella*, and *Prevotella* was significantly higher in the large intestine compared to the small intestine, whereas *Streptococcus* was more abundant in the small intestine than in the large intestine (Supplementary Figure 4). Except for menadione, which is mainly synthesized by *Bifidobacterium* and *Collinsella*, the remaining eight vitamins are synthesized by different bacterial genera in both the large and small intestines.

### 3.4 Genome catalog of vitamin de novo biosynthetic pathways

To further explore the ability of the feline gut microbiota to synthesize vitamin B and menaquinone at the genomic level, we selected 1,004 high-quality, non-redundant genomes (completeness ≥ 90%, contamination < 5%, and completeness - 5 × contamination ≥ 50) from a total of 7,324 genomes (Supplementary Table 4). These genomes were clustered by ANI similarity (> 99%). Genome annotation revealed that 782 of these genomes were predicted to be capable of synthesizing at least one vitamin B or menaquinone *de novo* (Figure 4A and Supplementary Table 5). The genome sizes of the 782 genomes capable of vitamin biosynthesis ranged from 1.04 Mbp to 6.20 Mbp (average 2.54 Mbp), with an average N50 length of 52,987.11 bp. The GC content varied from 16% to 73% (average 44.50%) (Figure 4B-C). Taxonomic assignment revealed that the two most abundant phyla were *Bacillota_A* (40.54%, 317 genomes) and *Actinomycetota* (20.84%, 163 genomes), followed by *Bacteroidota* (15.22%, 119 genomes), *Bacillota* (5.24%, 41 genomes), and *Bacillota_C* (4.60%, 36 genomes).

**Figure 4.**
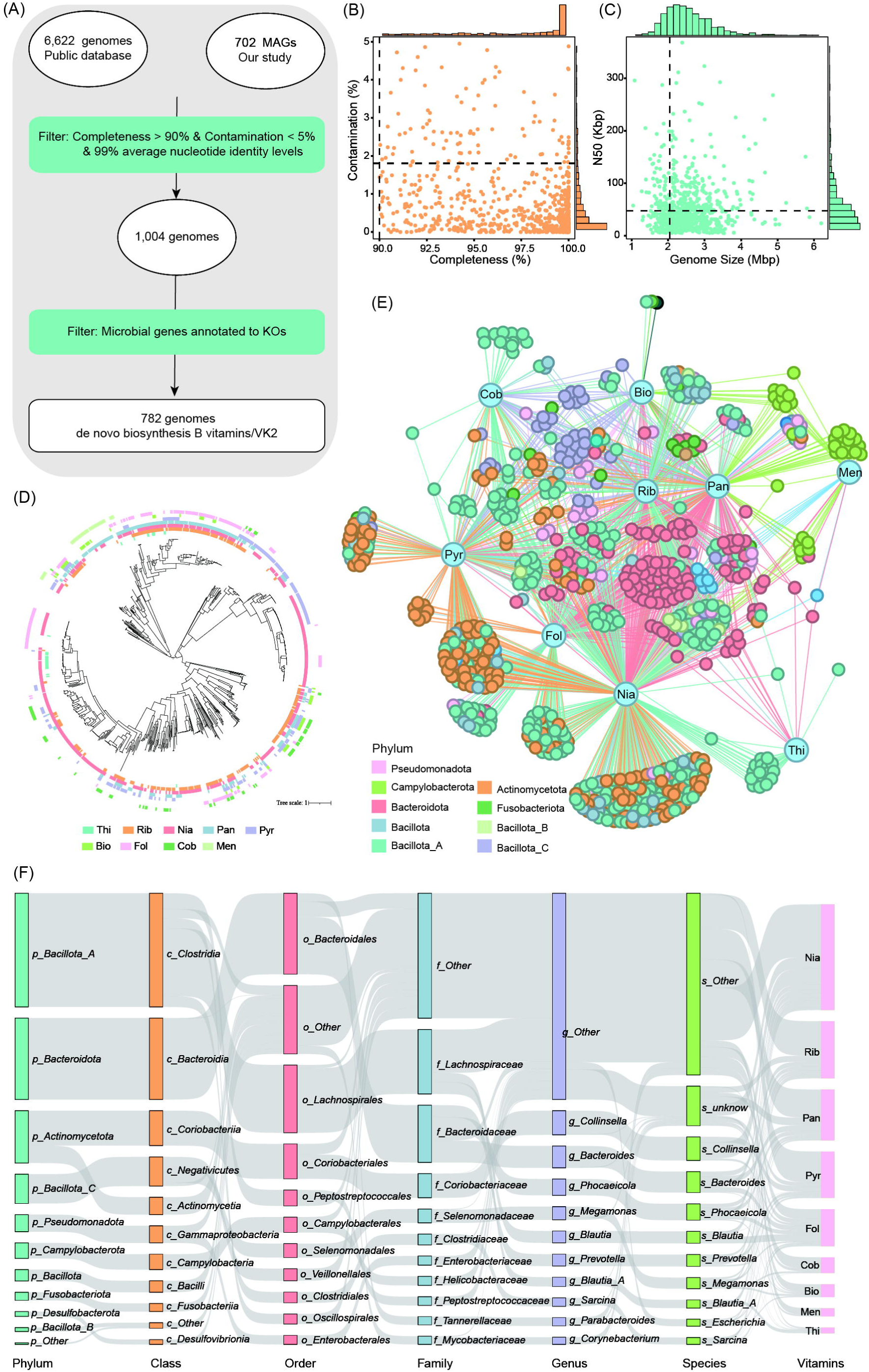
Comprehensive analysis of genomes synthesizing vitamins B and K_2_ across taxonomic levels. **(A)** Workflow for identifying genomes capable of synthesizing vitamins B and K_2_ (menaquinone) through a stepwise filtering process. Initially, 7,324 genomes were selected, filtered for completeness (>90%) and contamination (<5%), and further refined by removing redundancies (99% average nucleotide identity) and selecting those with annotated microbial genes linked to KEGG Orthologs (KOs). This resulted in 782 genomes capable of de novo biosynthesis of vitamins B and K_2_. **(B-C)** Genomic statistics for the 782 selected genomes, including genome size distribution (in megabase pairs, Mbp) and completeness versus contamination percentages. **(D)** Maximum-likelihood tree constructed for the 782 genomes, with clades color-coded based on their genome source. Outer-layer heatmaps indicate the presence (colored) or absence (blank) of specific vitamin synthesis capabilities. **(E)** Correlation network linking genomes and vitamin synthesis capabilities. Genomes are color-coded by taxonomic classification, and the relationships between genomes and vitamin synthesis capabilities are visualized. Abbreviations for the vitamins are as follows: thiamine (Thi), riboflavin (Rib), niacin (Nia), pantothenate (Pan), pyridoxine (Pyr), biotin (Bio), folate (Fol), cobalamin (Cob), and menaquinone (Men). **(F)** Sankey diagram illustrating the relationships between taxa (organized by taxonomic levels: phylum, class, order, family, and genus) and vitamin types. The diagram highlights taxonomic groups contributing to the synthesis of various vitamins.

We assessed the vitamin synthesis capabilities of the 782 genomes and found that 262 genomes could synthesize one vitamin, 459 genomes could synthesize two to four vitamins, and 60 genomes could synthesize five to six vitamins. No genome was capable of synthesizing more than eight vitamins *de novo*, with only one genome from the *Bacillota* phylum capable of synthesizing seven vitamins (excluding cobalamin and menaquinone) (Figure 4D). This suggests that most microorganisms likely acquire and synthesize vitamins through microbial interactions. The number of genomes classified for biosynthesis of each vitamin was as follows: Niacin: 557 genomes, riboflavin: 306 genomes, pantothenate: 273 genomes, pyridoxine: 249 genomes, folate: 198 genomes, cobalamin: 85 genomes, biotin: 68 genomes, menaquinone: 44 genomes, thiamine: 30 genomes (Figure 4E).

Our analysis of genome-based species annotations revealed that the synthesis of various vitamins in the cat intestine is primarily dependent on *Bacillota_A* and *Bacteroidota*. Specifically: riboflavin, pantothenate, folate, and thiamine are mainly synthesized by genomes from these two bacterial phyla, with *Bacillota_A* contributing 35.62%, 17.95%, 33.33%, and 66.67%, respectively, and *Bacteroidota* contributing 30.72%, 41.39%, 41.91%, and 30.00%, respectively. Niacin and pyridoxine are predominantly synthesized by genomes from *Bacillota_A* and *Actinomycetota*, with *Bacillota_A* contributing 43.39% and 27.71%, respectively, and *Actinomycetota* contributing 21.34% and 38.55%, respectively. Biotin is primarily synthesized by genomes from *Bacillota_C* (27.94%) and *Pseudomonadota* (16.18%). Menaquinone synthesis is primarily attributed to genomes from *Campylobacterota* (75.00%). Cobalamin synthesis is predominantly carried out by genomes from *Bacillota_A* and *Bacillota_C* (87.06%) (Figure 4F).

We also compared the genome catalogs for de novo vitamin synthesis across different animal species. The results revealed that the primary genomic contributors to vitamin biosynthesis in the cat intestinal microbiome are from *Bacillota_A* (40.54%) and *Actinomycetota* (20.84%). In contrast, in ruminants, the predominant contributors are genomes from *Bacteroidota* (43.28%) and *Bacillota* (38.25%). In the chicken intestine, *Bacteroidota* (28.21%) and *Bacillota_A* (25.66%) genomes are most abundant (Supplementary Figure 5). These findings highlight species-specific differences in the primary genomic contributors to vitamin biosynthesis.

### 3.5 *T. gondii* infection disrupts vitamin biosynthesis pathways

To explore the effects of *T. gondii* infection on the cat gut microbiota’s ability to synthesize vitamins, we reanalyzed metagenomic data collected before and after infection, focusing on genomic data related to vitamin biosynthesis pathways. Alpha diversity analysis was performed to assess the diversity of genes involved in vitamin B and menaquinone biosynthesis in both the small and large intestines. Alpha diversity analysis revealed that the diversity of genes involved in the biosynthesis of vitamin B and menaquinone increased significantly in the small intestine at 3 dpi, whereas no significant changes were observed in the large intestine (Figure 5A-B). In contrast, the diversity of *T. gondii*-related genes did not show significant changes in either the small or large intestine by the eighth dpi. PCoA based on Bray-Curtis distances further indicated that the gene composition of vitamin B and menaquinone biosynthetic pathways changed primarily in the small intestine after infection (Figure 5C). This suggests that the small intestine is more responsive to changes in vitamin biosynthesis following infection, while the large intestine remains relatively stable.

**Figure 5.**
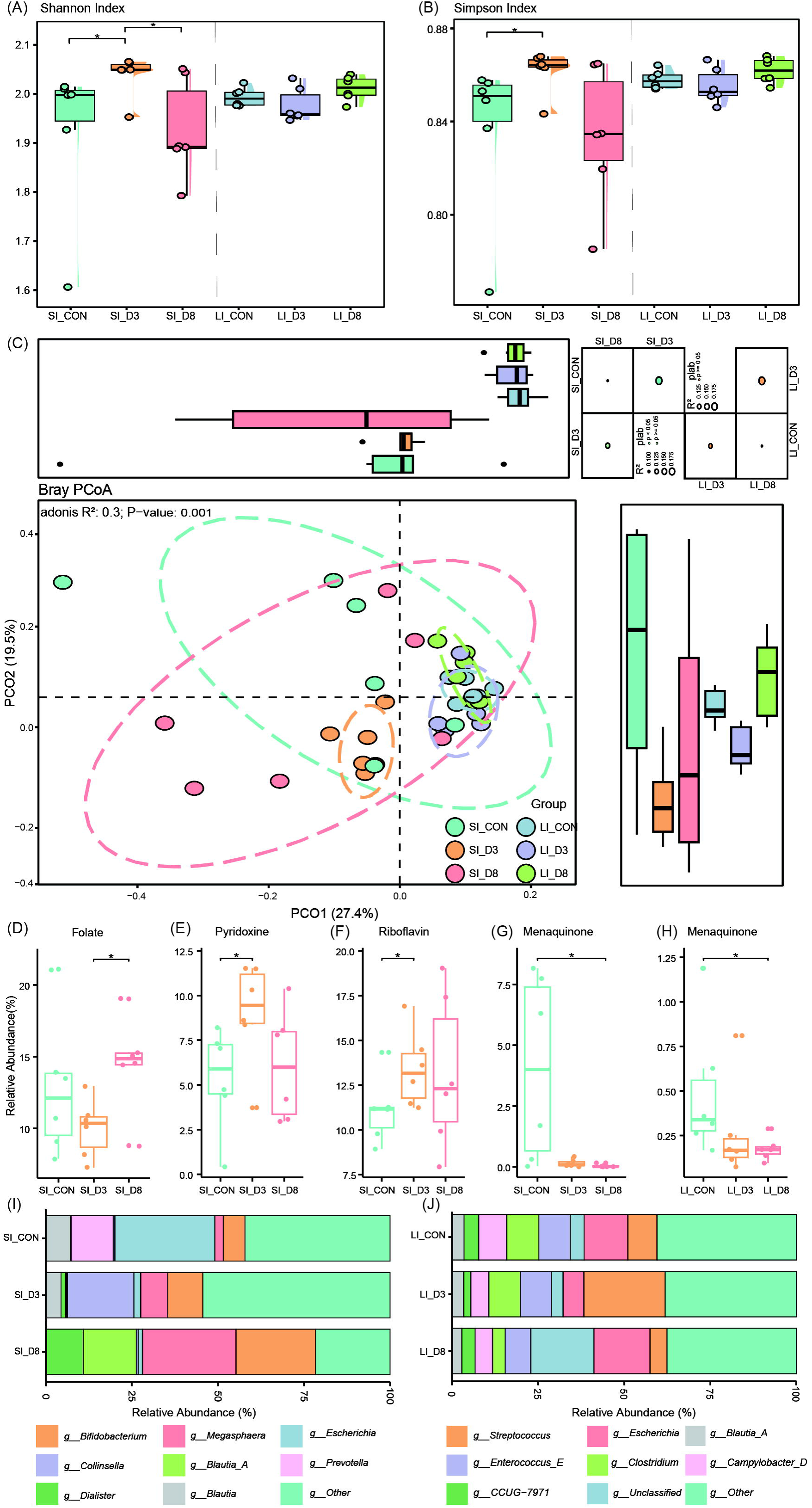
Impact of *Toxoplasma gondii* infection on microbial community structure and vitamin biosynthesis. **(A-B)** Raincloud plots show the distribution, probability density, and individual data points for the richness and Shannon index of microbial genomes involved in vitamin B and K_2_ biosynthesis across different experimental conditions. Boxplots summarize the distributions. Statistical significance was assessed using the Wilcoxon rank-sum test (* *p* < 0.05; ** *p* < 0.01; *** *p* < 0.001). **(C)** Bray-Curtis principal coordinate analysis (PCoA) visualizes microbial community structure at different stages of spawning (SI_CON, SI_D3, SI_D8, LI_CON, LI_D3, LI_D8). Ellipses represent 95% confidence intervals. Boxplots positioned above and to the right of the PCoA plot illustrate the distribution of sample scores for PCoA1 and PCoA2. The multivariate permutation test results in the upper right corner summarize pairwise comparisons for microbial composition in the SI and LI. **(D-G)** Boxplots show significant changes in the relative abundance of genes related to vitamin biosynthesis (Folate, Menaquinone, Pyridoxine, and Riboflavin) in the SI before and after *T. gondii* infection. **(H)** Boxplot illustrates significant changes in the relative abundance of genes related to vitamin biosynthesis (Menaquinone) in the LI across different experimental conditions. **(I-J)** Bar graphs show the phylum- and genus-level taxonomic distribution of genes related to vitamin B and K_2_ biosynthesis in the SI and LI. Taxonomic groups are color-coded, with relative abundances (%) presented for each condition. *Abbreviations:* SI, Small Intestine; LI, Large Intestine; CON, Control; D3, 3rd day after infection; D8, 8th day after infection. Statistical significance was determined using the Wilcoxon rank-sum test (* *p* < 0.05; ** *p* < 0.01; *** *p* < 0.001).

To further investigate the specific vitamins affected by *T. gondii* infection, we analyzed changes in the abundance of genes involved in synthesizing various vitamins across the two intestinal regions. Pyridoxine and Riboflavin synthesis in the small intestine significantly increased at 3 dpi (Figure 5D-E). Folate synthesis also significantly increased in the small intestine on the eighth day (Figure 5F). However, menaquinone synthesis decreased significantly in both the small and large intestines at 8 dpi (Figure 5H). No significant changes were observed in the synthesis of other vitamins.

To identify the microbial species contributing to these changes, we performed a taxonomic analysis of the vitamin biosynthetic genes. Among the top eight genera with the highest relative abundance, the following genera showed significant changes: *Blautia_A*, *Enterococcus_E,* and *Ligilactobacillus* exhibited a significant increase in the small intestine at 3 dpi. *Megasphaera* showed a significant increase in the large intestine at 3 dpi (Figure 5I-J and Supplementary Figure 6). These findings suggest that specific microbial genera are involved in modulating vitamin biosynthesis pathways in response to *T. gondii* infection.

### 3.6 *T. gondii* alters menaquinone synthesis pathways

Menaquinone was the only vitamin whose gene abundance exhibited significant changes in both the large and small intestines of cats following *T. gondii* infection. This observation prompted us to investigate how *T. gondii* disrupts menaquinone synthesis in the feline gut microbiota, as it could reveal critical insights into the infection’s broader metabolic consequences. Vitamin K_2_, a vital micronutrient for blood clotting, bone health, and immune regulation, emerges as a critical target for understanding the metabolic consequences of *T. gondii* infection. We analyzed the distribution of genes involved in the de novo synthesis of menaquinone. The results revealed that the feline intestinal microbiota synthesizes menaquinone indirectly via the 1,4-dihydroxy-6-naphthoate pathway. However, the key enzyme MqnB involved in this pathway was found at extremely low abundance. Following *T. gondii* infection, the relative abundance of MqnB further decreased in the small intestine on 3 dpi. Interestingly, 6-amino-6-deoxy-futalosine can bypass MqnB and directly synthesize dehydroxanthine-futalosine through K18284, promoting menaquinone synthesis even in the absence of MqnB. This suggests an alternative mechanism through which the microbiota can maintain menaquinone production during infection. Taxonomic analysis of the menaquinone biosynthesis genes revealed that these genes are predominantly found in *Actinomycetota* and *Campylobacterota* (Supplementary Figure 7). Specifically, the MqnX genes were primarily derived from *Actinomycetota*, while other essential genes, including MqnA, MqnE, MqnC, and MqnD, were largely from *Campylobacterota* (Figure 6).

**Figure 6.**
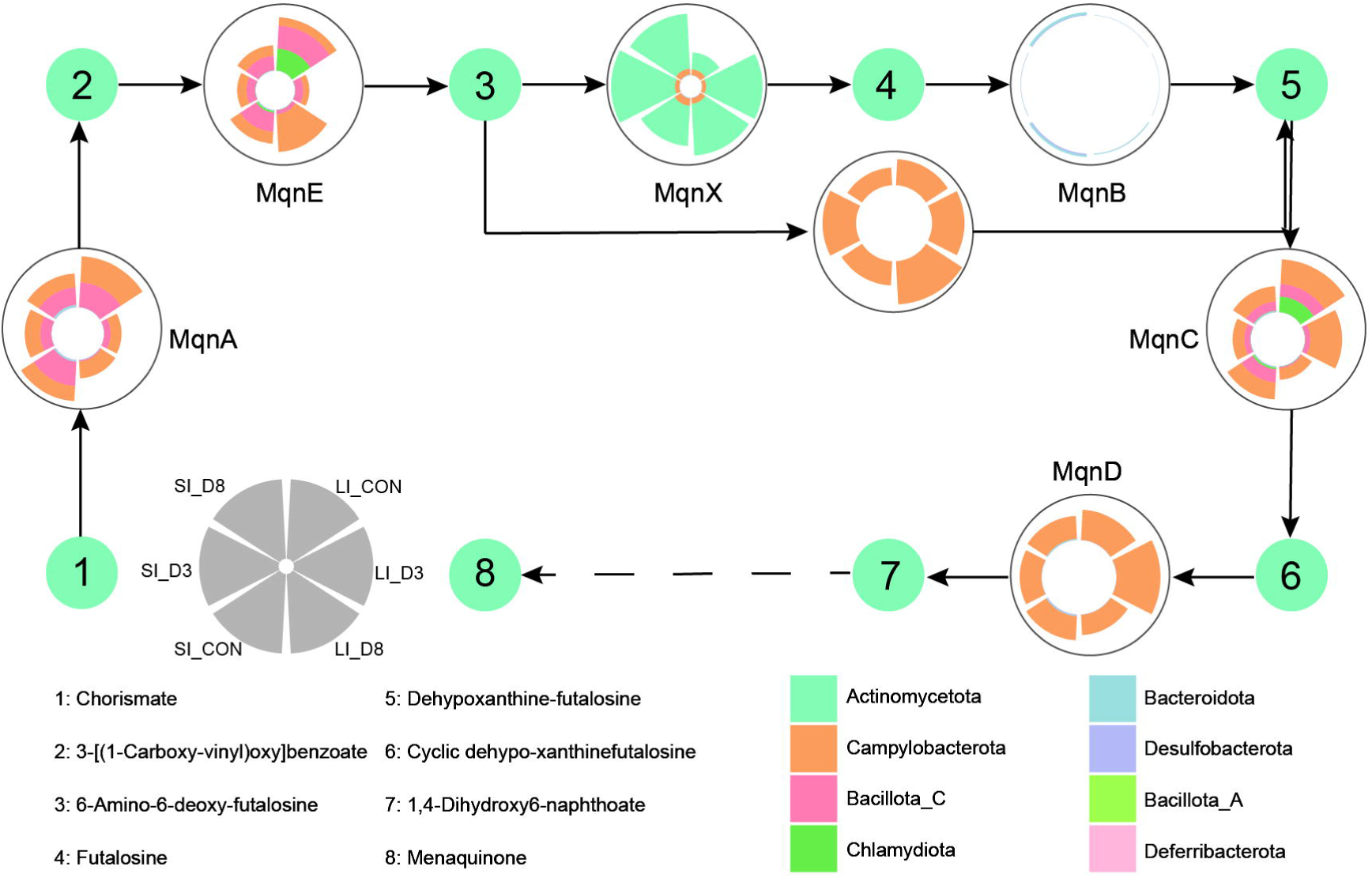
Distribution and taxonomic classification of genes involved in de novo biosynthesis of menaquinone across microbial phyla. Large circles represent distinct functional roles (e.g., MqnA, MqnB, MqnC, etc.) in the menaquinone biosynthetic pathway. Circular stacked bar charts within these circles illustrate the relative abundance and distribution of genes associated with these functional roles across various phylum-level classifications (e.g., *Actinomycetota*, *Campylobacterota*, and *Bacillota*). Each bar in the stacked chart is color-coded to indicate the contribution of specific microbial phyla to the biosynthetic genes. Small circles indicate the metabolites involved in each biosynthetic step, highlighting the biochemical intermediates produced during menaquinone synthesis. The figure emphasizes differences in gene abundance and phylogenetic diversity, underscoring the dominant role of certain phyla in menaquinone production.

## 4. Discussion

*Toxoplasma gondii* infection, a prevalent parasitic condition in cats, induces significant changes in the structure and function of the feline gut microbiome, which plays a critical role in synthesizing vitamins B and K_2_, essential for host metabolism and immunity (Bhattacharjee *et al*. 2017). As the primary site of *T. gondii* sexual reproduction, the feline intestine experiences inflammation and mucosal alterations that may disrupt microbial communities and their metabolic outputs, potentially affecting nutrient availability (Dubey 1997; Frenkel et al. 1970). To investigate these dynamics, we constructed a comprehensive gene catalog containing over 1.57 million genes from 1,553 non-redundant, high-quality metagenomes, enabling an in-depth analysis of vitamin biosynthesis pathways and their response to *T. gondii* infection.In addition, this study established a gene catalog containing more than 1.57 million genes, providing valuable insights into metabolism, host-microbe interactions and disease resistance.

In this study, we conducted an in-depth analysis of the biosynthetic pathways of vitamin B and K_2_ in the cat gut microbiome using the datasets mentioned above. Through this vitamin-related microbiome gene catalog, we demonstrated the spatial diversity of the biosynthetic abundance of vitamin B and K_2_ and the related microbial composition in the small and large intestines of cat gut microbes, consistent with previous studies (Deng *et al*. 2015; Handl *et al*. 2011), which is linked to its slower transit time, anaerobic conditions, and greater substrate availability (Donaldson *et al*. 2016a).

Vitamin biosynthesis pathways showed regional specialization: thiamine biosynthesis was predominantly mediated by the thI branch, while cobalamin synthesis involved both aerobic and anaerobic pathways. Folate biosynthesis was less prominent, suggesting dietary reliance, while menaquinone biosynthesis was low, further indicating dietary dependence for this vitamin. Cats likely rely on dietary sources for menaquinone, in contrast to herbivores or omnivores (Alessandri *et al*. 2020; Barry *et al*. 2012; Laflamme 2020; Deusch *et al*. 2014; Council. 2006). The functional specialization of the feline gut microbiome aligns with its obligate carnivore diet (Young et al. 2016), highlighting a higher need for microbially synthesized vitamins B like thiamine, riboflavin, and folate (LeBlanc et al. 2013a; Verbrugghe et al. 2013), while menaquinone and folate synthesis appears limited, reinforcing the dependency on dietary sources.

The study also revealed significant variability in vitamin synthesis capabilities among gut microorganisms. No genome could produce all nine essential vitamins, emphasizing the importance of microbial cross-feeding and cooperative interactions. This functional redundancy likely reflects evolutionary adaptation, ensuring sufficient vitamin production in an obligate carnivore host with inconsistent dietary sources (Alessandri et al. 2020) and highlighting the specialized roles of these taxa in gut vitamin biosynthesis.

Comparative analysis across species (ruminants, chickens, and cats) showed differences in microbial contributors to vitamin synthesis, reflecting host-specific adaptations. In cats, *Bacillota_A* and *Actinomycetota* predominate, aligning with the obligate carnivore diet, where microbial synthesis supplements dietary intake (Dodd *et al*. 2021; Hirakawa 1985.). This contrasts with ruminants and chickens, where *Bacteroidota* and *Bacillota* also play a major role. These findings underscore the role of microbial communities in supporting nutrient requirements, especially in diets with limited vitamin sources (Young et al. 2016; Studdert *et al*. 1991; Smith 1944).

*T. gondii* infection induced region-specific changes in gut microbial composition and gene diversity, with the small intestine showing greater sensitivity. At 3 dpi, the small intestine displayed a marked increase in genes associated with vitamin biosynthesis, particularly pyridoxine and riboflavin, both crucial for immune function and energy metabolism. By 8 dpi, folate biosynthesis also increased, likely reflecting microbial adaptation to heightened cellular turnover and immune activation. In contrast, menaquinone biosynthesis decreased significantly in both intestinal regions at 8 dpi, potentially due to the depletion of menaquinone-producing bacteria, such as *Campylobacterota*. These results suggest that the small intestine is more responsive to *T. gondii* infection, possibly due to its critical role in nutrient absorption and host-parasite interactions. As the site where *T. gondii* undergoes sexual reproduction in cats (Hutchison 1970; Frenkel 1970; Ferguson 1974), the small intestine may experience inflammation (Dias *et al*. 2014; Liesenfeld *et al*. 1996), immune activation (Benson et al. 2009), and pathological changes in the intestinal mucosa (Denkers 2009; Liesenfeld et al. 1996), which could influence the microbial community structure and its metabolic functions. The relative stability observed in the large intestine could reflect a more resilient microbiota, less affected by early infection-induced changes.

Taxonomic analysis revealed distinct microbial contributors to these changes. *Blautia_A*, *Enterococcus_E*, and *Ligilactobacillus* increased in the small intestine at 3 dpi, supporting B-vitamin production, while *Megasphaera* expanded in the large intestine, adapting to localized nutrient shifts and inflammation. These microbial shifts suggest a resilience mechanism to sustain vitamin production during infection, consistent with previous findings of *T. gondii*-induced changes in the metabolically active gut taxa (Hong et al. 2023). The observed changes are likely driven by several factors. Host immune responses, including inflammation (Bonnart et al. 2017; Benson et al. 2009) and altered gut conditions such as pH and oxygen levels, may favor certain bacterial taxa while suppressing others. *T. gondii* itself may compete with gut bacteria for nutrients like folate and pyridoxine, driving increased microbial biosynthesis (Massimine *et al*. 2005). Simultaneously, the infection reduces menaquinone-synthesizing bacteria while promoting the expansion of other taxa, altering metabolic outputs. Cross-feeding interactions among microbes may also help sustain vitamin production under the stress of infection (Magnusdottir *et al*. 2015). These findings emphasize the complex interplay between *T. gondii*, the gut microbiota, and the host. While the parasite alters the microbiota to meet its metabolic needs, these changes also influence host immune responses and resistance to infection (Partida-Rodriguez *et al*. 2017). Future research should explore how these dynamics benefit the host, the parasite, or both, and assess their long-term implications during infection.

*T. gondii* infection significantly disrupted menaquinone synthesis pathways, particularly in the small intestine. Reduced abundance of MqnB, a key enzyme in the canonical 1,4-dihydroxy-6-naphthoate pathway, highlights the vulnerability of this primary route as early as 3dpi. However, an alternative pathway via the 6-amino-6-deoxy-futalosine route is activated, enabling K18284 to bypass MqnB and directly synthesize menaquinone intermediates. This adaptation reflects microbial resilience under infection-induced stress, though the efficiency and overall contribution of this secondary pathway remain unclear.

Menaquinone biosynthesis in the feline gut microbiota is largely attributed to two phyla: *Actinomycetota* which predominantly harbor MqnX, facilitating alternative enzymatic mechanisms and *Campylobacterota* which provides key genes (MqnA, MqnE, MqnC, MqnD) essential for the canonical pathway. *T. gondii* infection may disrupt menaquinone synthesis by altering the abundance or activity of these microbial taxa, likely due to inflammation and nutrient imbalances in the gut. This disruption is consistent with the small intestine’s role as the primary site of host-parasite interactions and localized inflammation during infection (Egan *et al*. 2012; Elsheikha et al. 2021; Liesenfeld et al. 1996; Dias et al. 2014).

These disruptions reduce menaquinone production, potentially depriving the host of sufficient vitamin K_2_ — a nutrient critical for coagulation, bone health, and immune regulation. Vitamin K_2_ deficiencies resulting from *T. gondii* infection could have severe health consequences. For example, reduced synthesis of clotting factors increases the risk of bleeding disorders, as observed in vitamin K-deficient cats fed certain diets (Strieker et al. 1996) or malabsorption syndrome attributed to enteritis (Edwards et al. 1987). Diminished menaquinone availability may impair osteocalcin activation, compromising bone mineralization (Yamaguchi *et al*. 2011; Giordani *et al*. 2023). Deficiency could also intensify inflammation (Giordani et al. 2023) which may interfere with the host’s ability to control infection. However, due to the limited scope of our study, which examined only two time points, no definitive conclusions can be drawn regarding broader host health effects. These findings highlight the metabolic impact of *T. gondii* infection on microbial vitamin synthesis and underscore the need for future studies to quantify the role of classical and alternative pathways and their impact on feline physiology.

In summary, we compiled 1,553 non-redundant, medium- and high-quality feline gastrointestinal tract metagenomes through metagenomic assembly and public data collection. This enabled us to gain a comprehensive understanding of microbial-mediated vitamin B and K_2_ biosynthesis in the feline intestinal microbiome. We identified regional differences in vitamin synthesis between the small and large intestines, driven by distinct bacterial phyla. Notably, *T. gondii* infection initially enhances B vitamin production in the small intestine but subsequently reduces vitamin K_2_ (menaquinone) synthesis across both regions due to disrupted biosynthetic pathways, with only partial compensation from alternative routes. These findings suggest that *T. gondii* infection alters gut microbial metabolism, potentially impacting host nutrition and immunity. Further research is needed to elucidate the functional consequences of these microbial shifts on feline health and to explore targeted therapeutic interventions.

## Data Availability

Metagenomic sequencing data for this study have been deposited in the National Center for Biotechnology Information (NCBI) under the BioProject PRJNA1203944. All other data supporting the findings of this study are available in the paper and supplemental materials.

## Acknowledgements

Project support was provided by the National Natural Science Foundation of China (Grant No. 32172887), the NSFC-Yunnan Joint Fund (Grant No. U2202201), the National Key Research and Development Program of China (Grant Nos. 2021YFC2300800 and 2021YFC2300802), the Research Fund of Shanxi Province for Introduced High-level Leading Talents (Grant No. RFSXIHLT202101), and the Special Research Fund of Shanxi Agricultural University for High-level Talents (Grant No. 2021XG001). The study was supported by the Shandong Provincial Natural Science Foundation (No. ZR2025MS501), and the Key Laboratory of Veterinary Parasitology of Gansu Province Foundation (No. KLVPGP202502). We thank Shanghai Personal Biotechnology Co., Ltd., China. for technical assistance in metagenomic sequencing and preliminary analyses.

## Author Contributions

J.X.Z.: Writing – original draft, Formal analysis, Data curation, Investigation, Methodology, Visualization. X.Z.: Writing – original draft, Formal analysis, Data curation, Investigation, Methodology, Visualization. S.C.X.: Investigation, Methodology, Resources, Writing – review & editing. Y.H.L.: Investigation, Methodology, Resources, Writing – review & editing. Z.Z.: Investigation, Methodology, Resources, Writing – review & editing. Y.Q.G.: Formal analysis, Investigation, Writing – review & editing. L.Y.T.: Formal analysis, Investigation, Writing – review & editing. H.M.: Conceptualization, Project administration, Resources, Supervision, Writing – review & editing. X.X.Z.: Conceptualization, Formal analysis, Data curation, Project administration, Supervision, Funding acquisition, Writing – review & editing. All authors reviewed the manuscript.

