## Supplementary Information for "*Toxoplasma gondii* disrupts vitamin B and K2 biosynthetic pathways in feline gut microbiota: microbial adaptation"

### Supplementary Figures

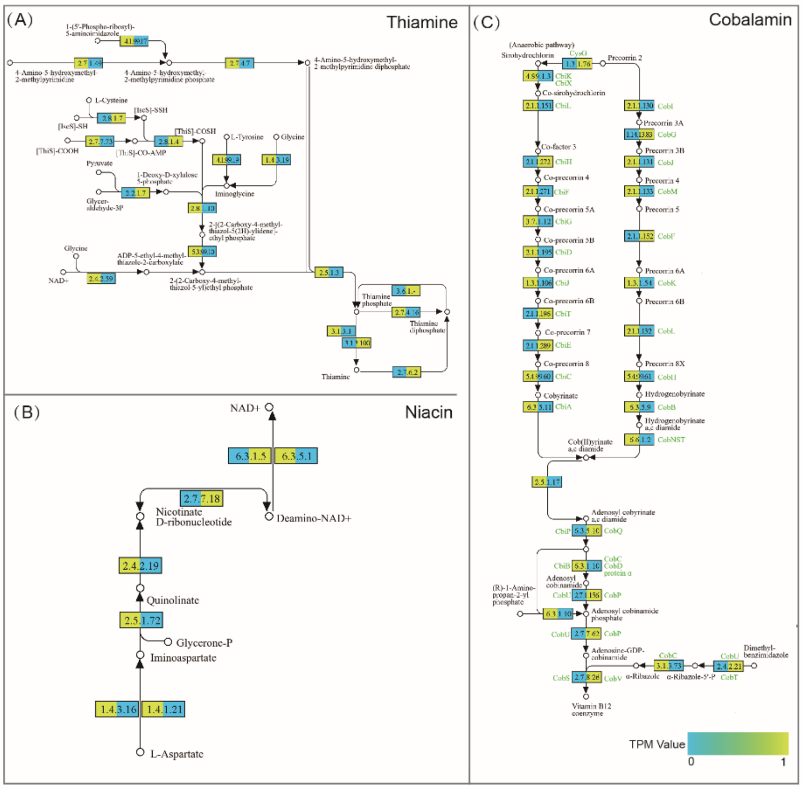


**Supplementary Figure 1. The biosynthesis pathways for thiamine (A), niacin (B), and cobalamin (C).** The rectangles and circles depict functional roles and metabolites, respectively. The rectangles are segmented into 2 sections, with different colors represent the values of different intestinal regions (TPM).

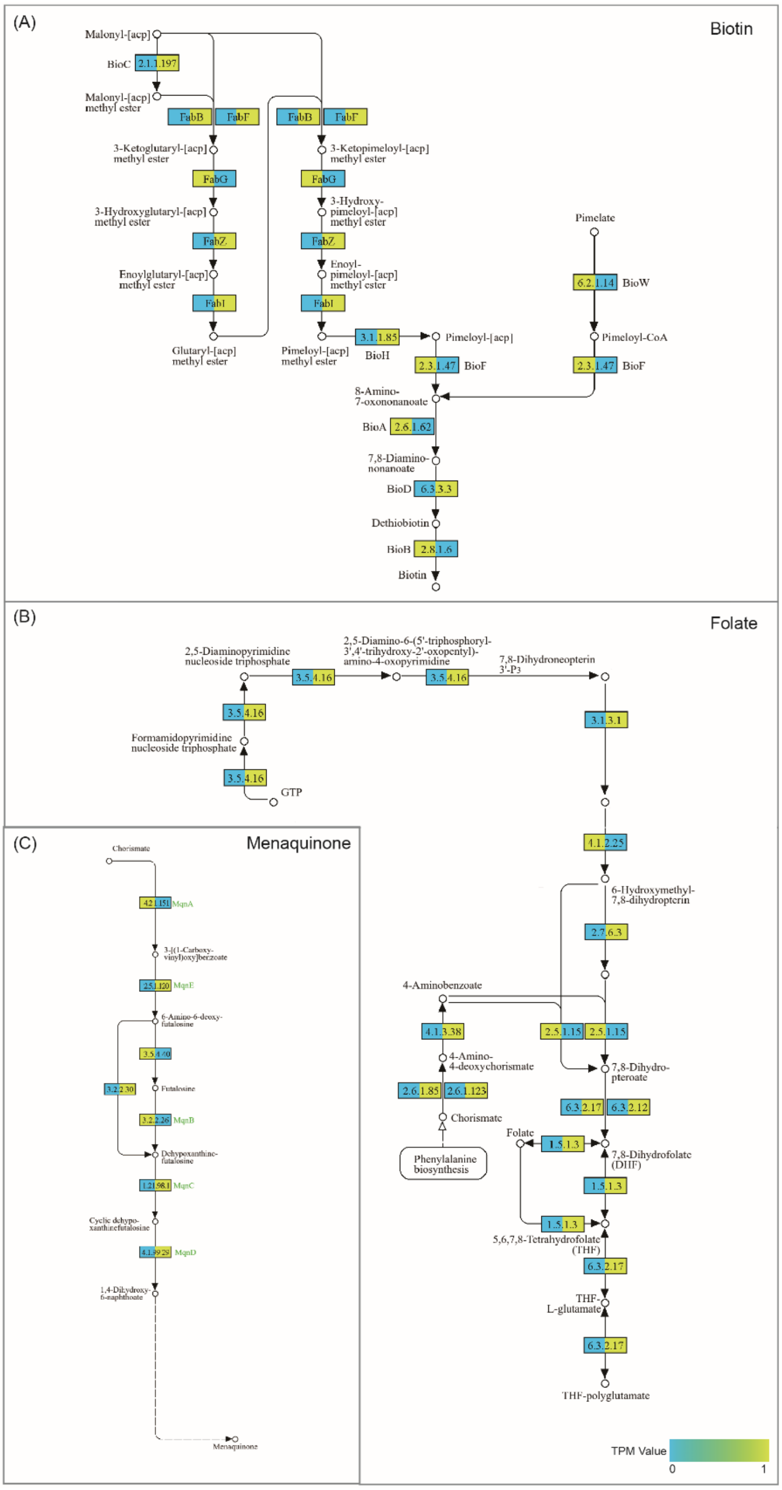


**Supplementary Figure 2. The pathways for (A) biotin, (B) folate, and (C) menaquinone biosynthesis.** The rectangles and circles depict functional roles and metabolites, respectively. The rectangles are segmented into two sections, with colors indicating TPM values across intestinal regions.

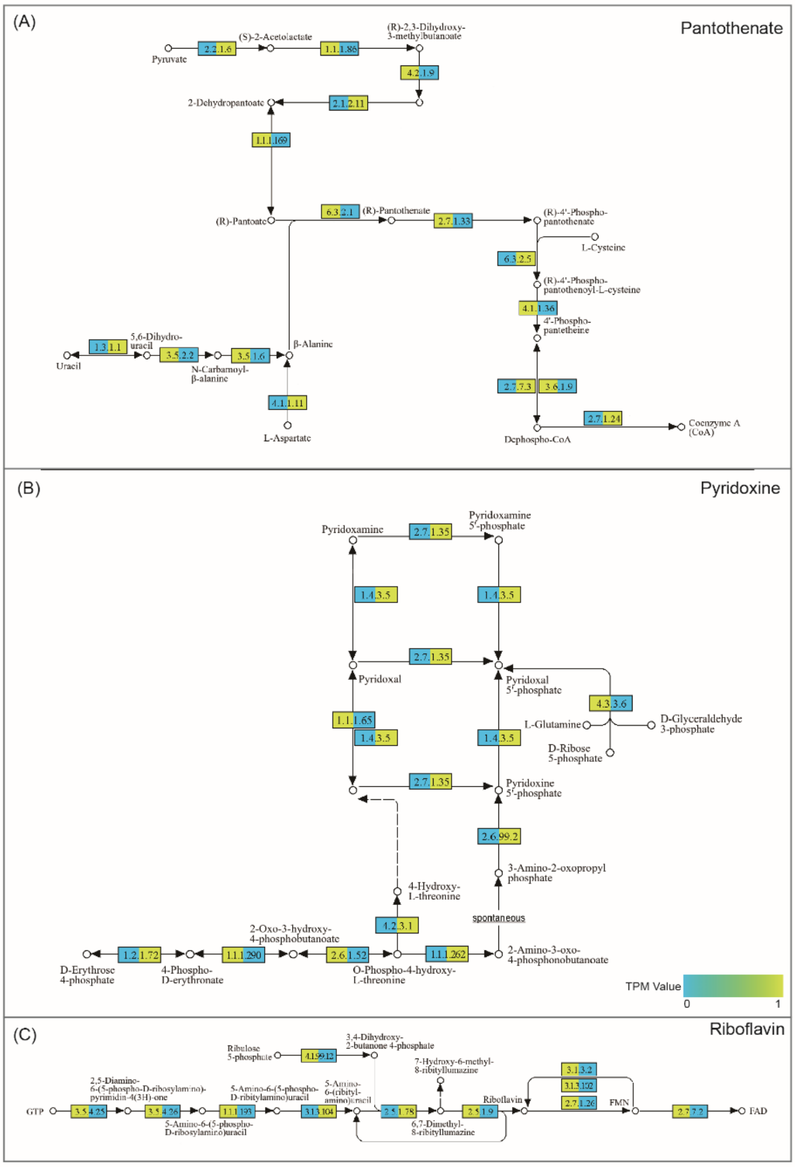


**Supplementary Figure 3. The (A) pantothenate, (B) pyridoxine, and (C) riboflavin biosynthesis pathways, with functional roles represented by rectangles and metabolites by circles.** The rectangles are segmented into two sections with different colors indicating TPM values across intestinal regions.

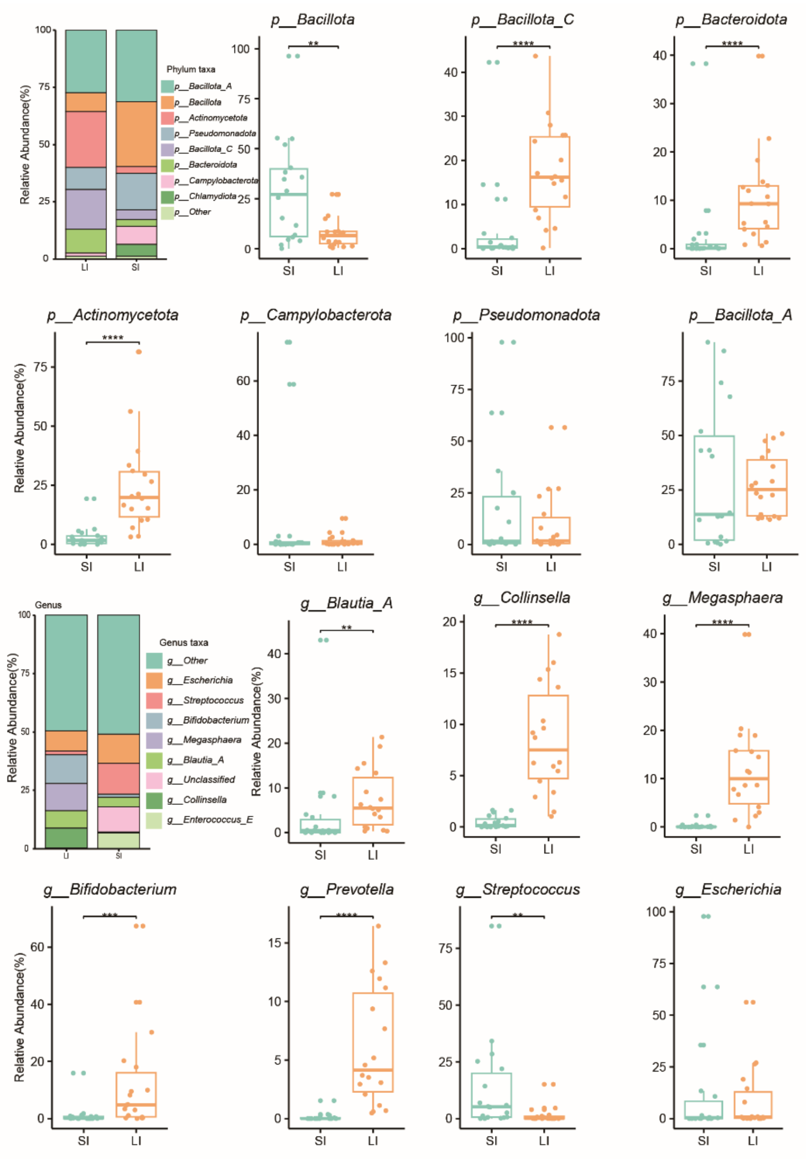


**Supplementary Figure 4. Distribution of vitamin biosynthesis-associated taxa across intestinal regions.** The bar graph presents the taxonomic distribution of genomes associated with vitamin B and K2 biosynthesis at the phylum and genus levels. The box plot compares the relative abundance of the top seven species between the small and large intestines. Statistical significance was determined using the Wilcoxon rank-sum test: * $p < 0.05$; ** $p < 0.01$; *** $p < 0.001$.

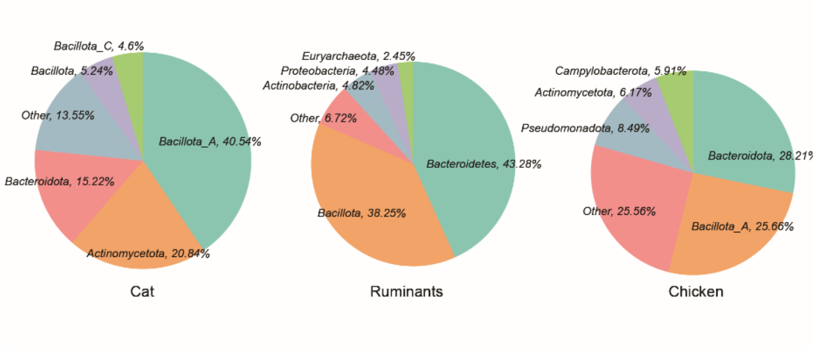


**Supplementary Figure 5. A pie chart displays the proportion of genomes capable of *de novo* vitamin synthesis at the phylum level across different animal hosts.**

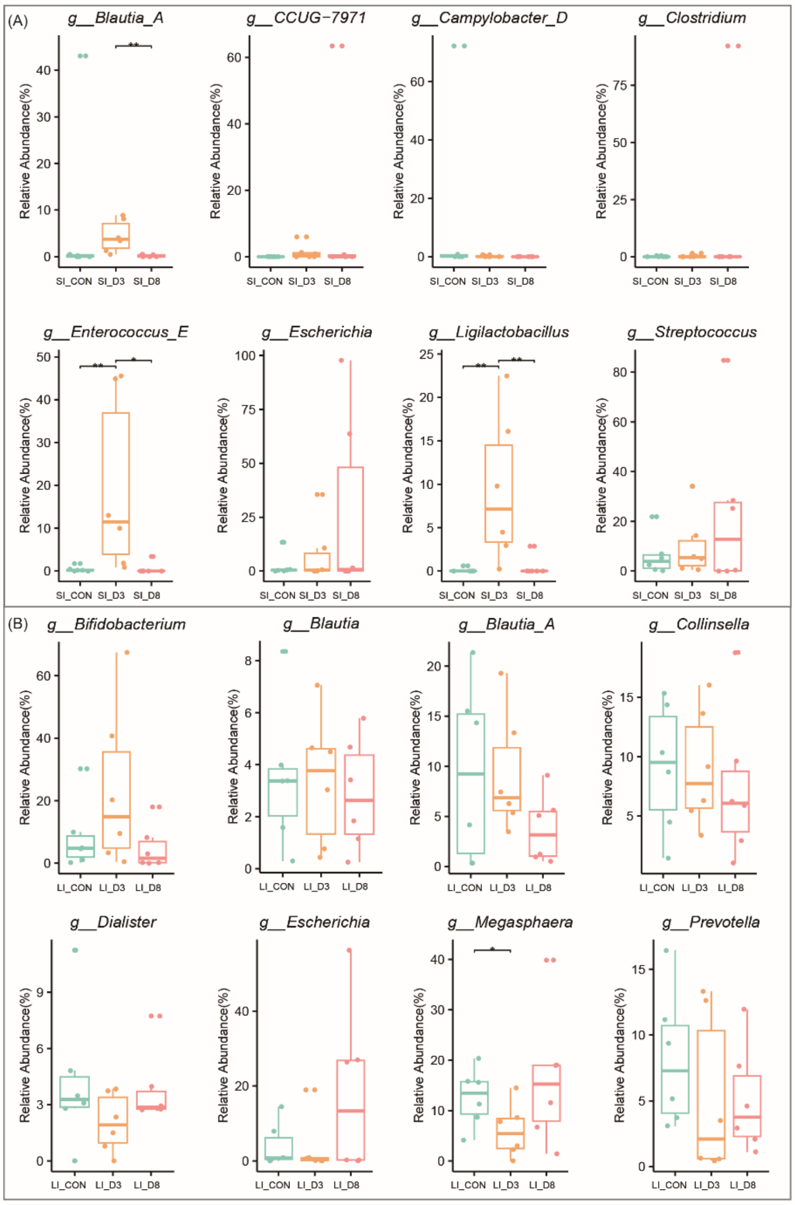


**Supplementary Figure 6. Impact of *Toxoplasma gondii* infection on the relative abundance of gut microbial genera. (A)** Box plot illustrating changes in the top eight most abundance genera in the small intestine before and after *T. gondii* infection. **(B)** Box plot illustrating changes in the top eight relative abundance genera in the large intestine before and after *T. gondii* infection. Wilcoxon rank sum test: * $p < 0.05$; ** $p < 0.01$; *** $p < 0.001$.

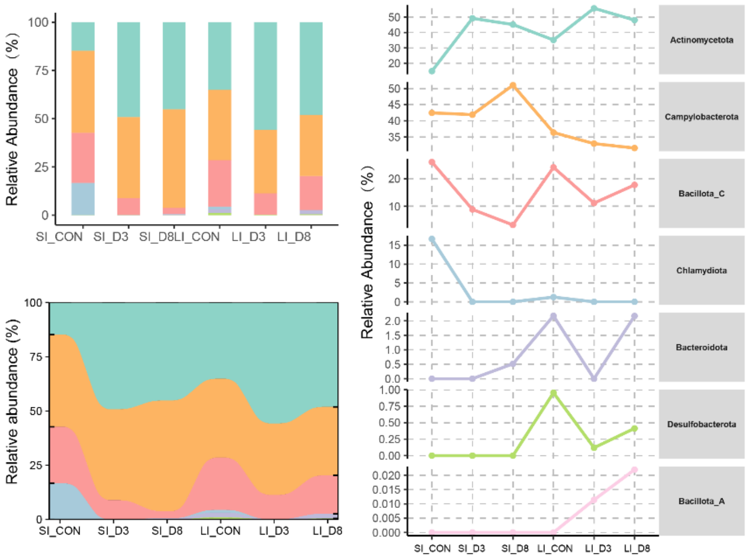


**Supplementary Figure 7. Relative abundance of menaquinone biosynthesis genes at the phylum level across the entire feline intestine, highlighting the top seven most abundant phyla.**

**Supplementary Table Legends**

**Supplementary Table 1:** Summary of 7,324 representative genomes, including their sources and quality assessment criteria.

**Supplementary Table 2:** Comprehensive details of genes involved in vitamin B and $K_2$ biosynthesis.

**Supplementary Table 3:** Phylum-level distribution of vitamin biosynthesis genes.

**Supplementary Table 4:** Taxonomic classification of 1,004 high-quality genomes.

**Supplementary Table 5:** Genomic statistics and vitamin synthesis capabilities of 782 selected genomes.
